# Machine learning–assisted selection of informative loci for strain-level phylogenetics of *Neisseria gonorrhoeae*

**DOI:** 10.1101/2025.10.02.679713

**Authors:** Elizaveta Kochubei, Serafim Dobrovolskii, Zlata Zenchenko, Mikhail Rayko

## Abstract

Epidemiological surveillance of *Neisseria gonorrhoeae* is hindered by the limitations of existing molecular typing methods, such as NG-MAST and MLST, which either suffer from excessive variability or insufficient resolution. In this study, we propose and evaluate a machine learning (ML) algorithm for the automated selection of a minimal set of informative genetic loci for accurate strain classification. Using a collection of 29 reference genomes of *N. gonorrhoeae*, we developed a pipeline that integrates Random Forest models and DNABERT embeddings to generate optimized gene panels. The results demonstrate that ML-selected panels substantially outperform traditional schemes, yielding markedly improved phylogenetic accuracy and branch support consistently above 90%. The proposed approach significantly reduces computational costs compared to whole-genome analysis and represents a promising resource-efficient tool for routine epidemiological monitoring, tracking transmission pathways, and identifying antibiotic-resistant strains.

## 1 Introduction

Metagenomic classifiers, thoroughly reviewed by Marić et al. (2024) [1], demonstrate satisfactory accuracy for species-level identification. However, when applied at the strain level, they face a number of fundamental limitations. K-mer–based classifiers, including Kraken2 [2] and Centrifuge [3], frequently yield false-positive assignments due to the high polymorphism of short oligonucleotides. Mapping-based approaches such as Minimap2 [4] and MetaMaps [5] are dependent on reference completeness and demand substantial computational resources, while a universal benchmark for long-read data has yet to be established.

Between 2023 and 2025, several tools emerged that partly mitigate these shortcomings. Demixer [6] combines Bayesian estimation of known clone proportions with de novo SNP-profile reconstruction, reducing reference bias, though it only performs reliably at coverage ≥ 20× and remains sensitive to recombination. MADRe [7] lowers false-positive detections through preliminary contig assembly and automatic database reduction, but at the cost of longer runtimes and the requirement of good coverage. Clasnip [8] introduces HMM-based typing of short amplicons via a web interface, increasing accessibility for screening, but remains entirely dependent on database updates and lacks support for mixed samples.

Alongside the publication of minMLST [9], where XGBoost was applied to select a minimal locus set from cgMLST schemes [10] while retaining adequate resolution (adjusted Rand index 0.4–0.93), marker-based metagenomic methods have been actively developed, such as StrainPhlAn [11]. Nevertheless, none of the currently available tools combine high strain-level resolution, moderate computational requirements, and full integration into routine diagnostics.

### 1.1 Specifics of *Neisseria gonorrhoeae*

*Neisseria gonorrhoeae* is characterized by extremely small intergenomic distances between circulating clones (≤0, 001 substitutions/site) and a high rate of recombination (r/m ≈2, 5) [12, 13, 14]. These features limit the effectiveness of the following:

- **MLST** [15] — the seven-housekeeping-gene scheme produces consistent but insufficiently detailed phylogenies;
- **NG-MAST** [16] — a two-gene system (*porB, tbpB*) improves resolution but generates polytomies that fail to capture resistance phenotypes or geographic distribution;
- **cgMLST** — leveraging 100–2000 core loci increases accuracy, yet remains resource-intensive and overlooks accessory genome variability;
- **Whole-genome analysis** [17] — provides maximal phylogenetic resolution but demands substantial computational capacity, high sequencing costs, and expert interpretation.

Marić et al. demonstrated that the absence of a single species from a reference database introduces ≥20% error in abundance estimation; at the strain level this is equivalent to missing a clinically significant clone. Furthermore, k-mer classifiers tend to generate artificial “strains”, while resource-heavy mapping approaches are difficult to implement in laboratories with limited computational infrastructure.

### 1.2 Rationale

Effective epidemiological surveillance of *N. gonorrhoeae* a specialized, resource-efficient pipeline with a curated database of gonococcal genomes, a targeted subset of orthologs, and integrated resistance gene analysis.

Our project proposes an algorithm for automated selection of a minimal set of informative orthogroups (not restricted to core genes), subsequent assessment of phylogenetic tree topological stability, and simultaneous identification of antimicrobial resistance determinants. This approach preserves strain-level resolution, reduces computational costs, and enhances the method’s practical applicability.

The significance of this study stems from the persistently high global prevalence of gonococcal infection, caused by *N. gonorrhoeae*, which remains a major public health challenge. Particularly concerning is the rapid emergence of antimicrobial resistance in gonococcus, necessitating continuous and effective epidemiological monitoring.

The proposed approach has considerable practical utility. The developed algorithm can optimize PCR-based diagnostics by enabling the design of more accurate and cost-effective PCR assays targeting key epidemiological markers, including resistance genes. It is also critical to note that horizontal gene transfer plays a central role in the rapid spread of resistance determinants among *N. gonorrhoeae* populations, making their surveillance essential for effective disease control. We anticipate that our pipeline will facilitate timely identification of novel strains, track their dissemination pathways, and predict resistance development, thereby significantly strengthening infection control efforts.

## 2 Materials and Methods

### 2.1 Genomic data collection

We used the WHO 2024 reference panel of 29 *N. gonorrhoeae* strains [18], which covers major resistance phenotypes and genotypes, including clone FC428 (*penA*-60.001) and mosaic variants of the MtrRCDE efflux system. Genomes were sequenced using PacBio or Nanopore platforms with Illumina polishing, ensuring high assembly quality.

Key advantages of this dataset include:

- **Controlled diversity:** strains represent globally distributed clones, including extensively resistant lineages.
- **Comprehensive annotation:** phenotypic susceptibility profiles and genetic markers (e.g., *blaTEM, tetM*, 23S rRNA A2059G/C2611T, *gyrA* S91F/D95G) are available.
- **Nonredundancy:** unlike open repositories, the collection excludes duplicates and technical artifacts.
- **Standardization:** strains are employed in GASP and EGASP programs, facilitating cross-study comparability.

The dataset includes 15 newly designated (2024) and 14 historical reference strains, ensuring both temporal and geographic representativeness. Metadata comprise isolation site and year, MLST, NG-MAST, and NG-STAR profiles, allowing comparison with conventional typing methods.

Complete and chromosomal assemblies were retrieved from RefSeq [19] using *ncbi* − *genome* − *download* (level “complete/chromosome”; formats: GenBank + CDS FASTA). From the global pool, the 29 WHO-2024 isolates were selected; duplicates of historical strains (marked «_2024») were removed using regular-expression filtering. For each assembly, BioSample identifiers and strain names were parsed from *assembly*_*summary* files and incorporated into CDS headers using a custom Biopython script.

### 2.2 Ortholog detection and sequence alignment

Single-copy orthogroups were identified from downloaded CDS sequences using OrthoFinder (v2.5.5) [20] in DNA mode (−− *dna*). The resulting orthogroups were used for downstream phylogenetic analysis. Within each orthogroup, multiple sequence alignments were performed with MAFFT (v7.525) [21] using the −−*auto* setting. For each alignment, Shannon entropy and additional variability metrics were calculated to quantify locus diversity.

### 2.3 Reference and alternative phylogenetic reconstructions

#### Reference tree

To benchmark typing accuracy, a reference phylogeny was reconstructed from all single-copy orthogroups identified by OrthoFinder (1,776 genes). Orthogroup sequences were concatenated per strain, and trees were inferred using IQ-TREE2 [22] under maximum likelihood with automatic model selection (MFP, HKY+F+G4) and 1,000 ultrafast bootstrap (UFBoot) replicates.

#### Stepwise selection of variable loci

Ranked gene lists were used to build concatenated subsets of 1… N loci. For each subset, maximum-likelihood trees were reconstructed as above. Topological stability was evaluated by normalized Robinson–Foulds (RF) distance relative to the reference tree, complemented by mean bootstrap support.

#### Robinson–Foulds (RF) metric

The Robinson–Foulds distance (RF) measures the symmetric difference between two unrooted trees, expressed as the number of bipartitions (“splits”) that are present in only one of the compared trees. For two identical topologies, *RF* = 0; if clades do not overlap at all, RF reaches its maximum, calculated as 1:

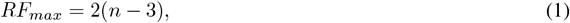

where *n* s the number of leaves. For *n* = 29 strains, the maximum RF equals 52. In this study, we additionally used the *normalizedRF* distance:

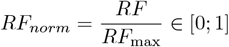

Values of *RF*_*norm*_ ≤ 0.20 were considered to represent “good” topological concordance, while *RF*_*norm*_ ≤ 0.10 was treated as virtually indistinguishable from the reference topology.

#### Median-variability controls

In addition to the subsets of the N most variable loci, control panels were assembled from orthogroups positioned near the median Shannon entropy value. Specifically, each set included nine loci: four immediately below and five immediately above the median [23].

This design was intended to test whether loci of only “moderate” informativeness can maintain phylogenetic signal, as opposed to hypervariable markers that may be biased by recombination or saturation.

For each control set, sequences were concatenated, aligned, and phylogenetic trees were reconstructed with IQ-TREE (1,000 ultrafast bootstrap replicates). Normalized Robinson–Foulds distances relative to the reference topology were calculated, and mean bootstrap support was recorded as an additional stability metric.

The comparative results for top-variable, median-variability, and ML-selected panels are presented side by side in the Results section.

### 2.4 ML-based gene selection(data preparation)

#### Generation of training data

From the 200 most variable orthogroups, random combinations of two, three, and four genes were generated, with three independent replicates per gene. For each combination, concatenated alignments were built, and phylogenetic trees were reconstructed using IQ-TREE. The target variable was the normalized RF distance from the reference phylogeny, computed with ETE3. Mean bootstrap support of the tree served as an independent reliability metric.

#### Feature engineering

For each orthogroup, eight quantitative descriptors were computed:

- *Mean*_*Edit*_*Distance* - the average number of operations (substitution, insertion, deletion) required to transform one allele sequence into another. Higher values indicate greater divergence between alleles.
- *TopK*_*Coverage* — the proportion of strains represented by the K most frequent alleles. For example, if the three most common alleles occur in 23
- *Mean*_*Normalized*_*Levenshtein* — the average pairwise Levenshtein distance between sequences, normalized by sequence length (range 0–1). A value of 0 indicates identity; 1 indicates maximal divergence.
- *SNP* _*Density* — the percentage of polymorphic positions (single-nucleotide substitutions) relative to the total sequence length.
- *GapFraction* — the fraction of positions in the multiple alignment represented by gaps (insertions/deletions), relative to alignment length.
- *LengthV ar* — the variance of allele lengths within the locus. Higher values reflect greater heterogeneity in sequence length.
- *UniqueAlleles*— the total count of distinct allelic sequences observed in the dataset for that locus. Larger counts indicate higher allelic richness.

These features were later used to train machine learning models aiming to minimize *RF* distance between reduced-locus trees and the reference phylogeny.

#### Embedding approach — DNABERT

For the embedding-based approach, all possible pairs were generated from the 200 most variable orthogroups, and multiple sequence alignments were concatenated for each pair. If a sequence exceeded 512 nucleotides, it was split into tokens of 512 bases (as required by the pretrained DNABERT model). Each fragment was passed through DNABERT [24], after which averaging was applied across all k-mers and across all fragments, yielding a single 768-dimensional embedding vector for each orthogroup pair.

### 2.5 ML-based gene selection (model training)

#### Processing of gene combinations

During model training, we used data obtained from random orthogroup combinations. For each combination of two, three, or four genes (with three independent replicates), concatenated alignments were generated, and phylogenetic trees were built in IQ-TREE. The target variable was the normalized Robinson–Foulds (*RF*) distance between the tree of the combination and the reference phylogeny, calculated with ETE3. As an additional reliability metric, the mean bootstrap support of each tree was considered.

For each orthogroup, the eight structural–evolutionary features described above (edit distance, allelic diversity, SNP density, gap fraction, etc.) were precomputed. For every combination of genes, these feature vectors were aggregated to form a single descriptor of the combination. A Random Forest regression model was trained to predict tree accuracy from these aggregated features. Data were split into training and test sets (e.g., 80/20), and fivefold cross-validation was applied. Each example was represented by the aggregated feature vector of the loci included, while the target variable was the normalized RF distance from the reference tree.

Based on feature importance and Random Forest predictions, we compiled a ranked list of individual genes statistically associated with low *RF* values (i.e., higher concordance with the reference phylogeny).

In a parallel analysis, the same model was trained on all possible orthogroup pairs drawn from the 200-variable set. For each pair, combined feature vectors were calculated, and the model was trained to predict RF distances. Pairs with minimal predicted RF were then used to reconstruct trees, which served to verify the accuracy of the ranking. Thus, the ML approach enabled identification of both individual loci and gene pairs with the greatest potential for accurate phylogenetic reconstruction.

#### Processing of embeddings for orthogroup pairs

For each of the C(200, 2) orthogroup pairs, embeddings were generated using DNABERT. Instead of training a model, we computed Euclidean distances between all embedding vectors and selected pairs with maximal separation (i.e., those most distinct in feature space). These pairs were then used for concatenation, tree building in IQ-TREE, and calculation of normalized RF distances relative to the reference tree. The rationale was that maximal distance in embedding space reflects complementary locus information, leading to reduced RF values even without additional ML prediction.

### 2.6 Validation of ML panels and Comparison with classical MLST schemes

The loci selected by the ML approach were concatenated according to the scheme “1… 10 genes.” For each panel size, a phylogenetic tree was reconstructed in IQ-TREE with 1,000 ultrafast bootstrap (UFBoot) replicates.

Topological accuracy was evaluated by the normalized Robinson–Foulds (RF) distance relative to the reference tree. Tree stability was additionally assessed by mean bootstrap support across branches.

To provide a direct benchmark, two historically used typing schemes were reimplemented and compared with the experimental panels:

#### NG −MAST

For each strain, CDS fragments of porB and tbpB were extracted, concatenated, aligned with MAFFT, and used to build a two-gene maximum-likelihood tree. The resulting topology was compared to the reference phylogeny using the Robinson–Foulds distance.

#### *pubMLST* (seven housekeeping genes)

Internal fragments of *abcZ, adk, aroE, fumC, gdh, pdhC*, and *pgm* were extracted according to the original MLST scheme. Each locus was aligned separately, then concatenated into a seven-gene panel. Trees were reconstructed using maximum likelihood with the same substitution model and 1,000 UFBoot replicates. Topologies were again compared to the reference using the Robinson–Foulds metric.

Thus, both conventional schemes (the two-gene NG-MAST and the seven-gene pubMLST) were evaluated alongside variable-locus panels and ML-selected panels, providing a unified comparative framework for all typing methods.

A fully reproducible implementation of all described steps — from genome retrieval to calculation of phylogenetic metrics and machine learning–based locus selection — has been deposited in an open GitHub repository [25], which includes all scripts, environment configurations, and example command lines for running the pipeline.

### 2.7 Antimicrobial resistance gene (ARG) analysis

To assess the clinical significance of the phylogenetic clusters, we performed parallel screening of antimicrobial resistance genes using the Comprehensive Antibiotic Resistance Database (CARD). For each of the 29 assemblies, the Resistance Gene Identifier (RGI) module was applied in “perfect” + “strict” modes, enabling identification of ARGs through sequence comparison with the curated CARD database.

Output tables from RGI were aggregated and processed in Python (v 3.9.12). using the libraries pandas, seaborn, and matplotlib.pyplot. Strain names were extracted from file identifiers, and antibiotic annotations were cleaned and normalized according to CARD ontology, including disambiguation of multiple antibiotics listed in a single record.

Resistance mechanisms (e.g.,(*β*-lactamase), efflux, target modification) were categorized following the CARD classification framework. For further analysis, the RGI text files were directly imported.

## 3 Results

### 3.1 ML-based approach

Cross-validation of the Random Forest model yielded a mean absolute error (MAE) of 0.042 RF units, indicating that the trained model reliably predicts topological divergence (Robinson–Foulds, RF) between trees constructed from arbitrary locus combinations and the reference phylogeny.

To quantify the contribution of algorithmic selection, we compared phylogenetic trees reconstructed from “top-variable” genes with those assembled from ML-predicted panels. The results are summarized in Table1 1 1.

As shown in Table 1, for panels of 2–4 genes, the ML-selected loci achieved 7–9 % lower normalized RF distances (*RF*_*norm*_ = 0.769) while maintaining comparable or higher mean bootstrap values. At seven loci, both strategies converged on the same RF distance, but the ML panel exhibited the highest branch support (91 %). The most pronounced improvement was observed for 10 loci, where the ML panel reduced *RF*_*norm*_ to 0.654 — 0.077 points lower than the corresponding variable-gene panel.

**Table 1:**
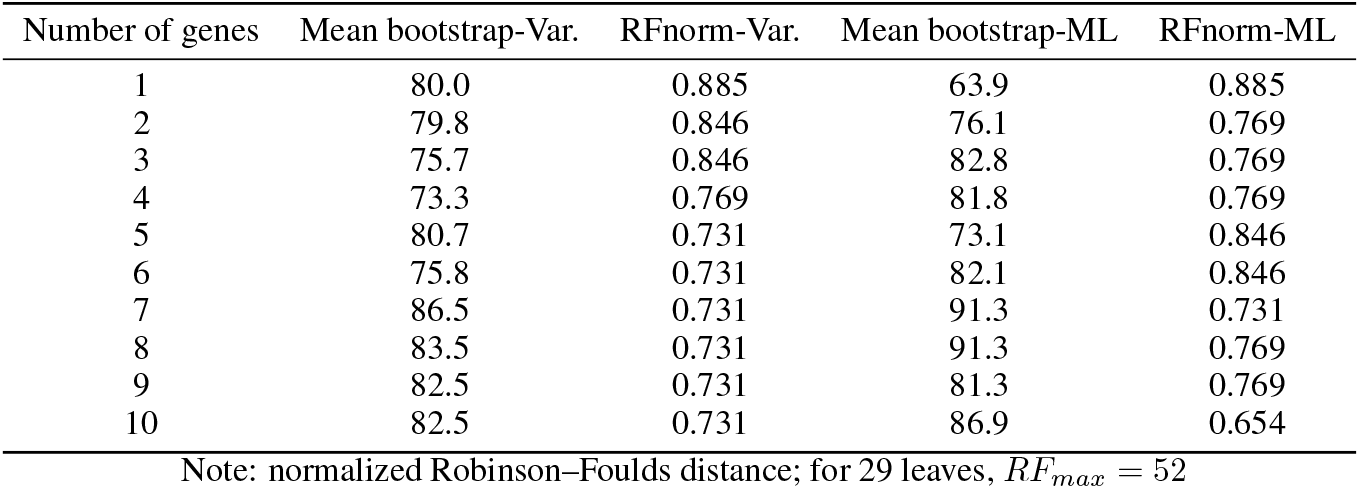
Comparison of “top-variable” and ML-predicted panels.

### 3.2 Predicted gene pairs

In the following results, orthogroups are designated as OG + number (e.g., OG0000159). Each orthogroup identified in our analysis was mapped to protein sequence identifiers from the RefSeq database, all of which carry the *WP* _. These *WP* _*ID* epresent reference records of prokaryotic proteins: when identical or nearly identical amino acid sequences are shared across strains or species, they are linked to the same *WP* _ entry.

Thus, a single orthogroup corresponds to a cluster of homologous proteins and may include multiple *WP* _*ID*, reflecting different alleles of the same gene or minor sequence-length variations across genomes. The presence of multiple *WP* _*ID* within one orthogroup indicates either allelic diversity or small variants that still fall into a common homology group. Consequently, the *WP* _*ID* for each orthogroup documents the specific protein sequences grouped into a single evolutionarily supported cluster. The complete mapping for each orthogroup is summarized in Table 6.

The Random Forest model identified several orthogroup combinations associated with improved topological accuracy. For each predicted pair, mean bootstrap support and normalized RF distance relative to the reference tree are summarized in Table 2.

**Table 2:** Results of predicted orthogroup pairs.

| Orthogroup pair | Mean bootstrap | RF |
| --- | --- | --- |
| OG0001093_OG0001710 | 63.261 | 0.769 |
| OG0001264_OG0001710 | 69.433 | 0.807 |
| OG0001093_OG0000461 | 63.504 | 0.769 |
| OG0001093_OG0001424 | 64.392 | 0.807 |
| OG0001710_OG0000625 | 72.213 | 0.961 |
| OG0001093_OG0000705 | 74.521 | 0.769 |

The Random Forest model predicted several gene pairs with favorable performance: normalized RF ≈0.769 −0.807 combined with mean bootstrap support of 63–74%. The best bootstrap value was observed for the pair OG0001093_OG0000705 (74.5%), though its RF remained at 0.769. This indicates a balance between improved tree stability and close correspondence to the reference phylogeny.

### 3.3 Results of distant orthogroup pairs by embeddings

The DNABERT-based embedding approach revealed orthogroup pairs with maximal separation in feature space. For each pair, mean bootstrap support and normalized RF distance relative to the reference phylogeny are reported in Table 3.

**Table 3:** Results of the most distant orthogroup pairs by embeddings.

| Orthogroup pair | Mean bootstrap | RF |
| --- | --- | --- |
| OG0000200_OG0001901 | 81.263 | 0.846 |
| OG0001547_OG0001304 | 71.792 | 0.884 |
| OG0001262_OG0001277 | 64.044 | 0.923 |
| OG0001202_OG0001438 | 64.155 | 0.961 |
| OG0001845_OG0000159 | 88.956 | 0.884 |
| OG0000200_OG0001901 | 81.261 | 0.844 |

The embedding approach allowed identification of orthogroup pairs with maximal divergence in feature space, which in most cases ensured higher bootstrap support (up to ≈89%) with moderate RF values (0.846–0.961). Of particular note is the pair OG0000200_OG0001901, which yielded two nearly identical results (RF = 0.846 and 0.844), demonstrating method stability across concatenated and fragmented alignments.

### 3.4 Comparison with classical MLST schemes

To evaluate how well established routine methods reproduce whole-genome phylogeny, the same metrics were calculated for NG-MAST and the seven-gene pubMLST scheme. Results are presented in Table 4).

**Table 4:** Topological accuracy of traditional typing schemes.

| Scheme | Number of genes | Mean bootstrap | RF | RFnorm |
| --- | --- | --- | --- | --- |
| NG-MAST (*porB*, *tbpB*) | 2 | 72 | 48 | 0.923 |
| pubMLST (7 genes) | 7 | 64 | 42 | 0.808 |

The two-gene NG-MAST panel reproduces only ≈8%% of the reference bipartitions, despite moderate bootstrap support (72%); its normalized RF of 0.923 indicates almost complete discordance with global topology. The seven-gene pubMLST improves slightly (*RF*_*norm*_ = 0.808; 12% concordance) but remains substantially worse than any experimental panel of ≥ 3 genes (see Table 1).

Collectively, these results confirm that both traditional schemes are insufficient for accurate intraspecies phylogenetic reconstruction of *N. gonorrhoeae*, and perform markedly worse than the ML-selected panels, which achieve nearly twofold lower RF distances with comparable or higher branch support.

### 3.5 Control panels of “median” variability

To evaluate whether loci of only moderate variability can preserve phylogenetic signal, we assembled control panels from orthogroups located near the median Shannon entropy value. The results of these median-variability panels are shown in Table 5.

**Table 5:** Evaluation of median-variability panels.

| Number of genes | RF | RFnorm | Mean bootstrap |
| --- | --- | --- | --- |
| 1 | 46 | 0.885 | 70.8 |
| 2 | 36 | 0.692 | 66.2 |
| 3 | 44 | 0.846 | 57.3 |
| 4 | 32 | 0.615 | 62.0 |
| 5 | 38 | 0.731 | 63.7 |
| 6 | 42 | 0.808 | 61.9 |
| 7 | 36 | 0.692 | 67.3 |
| 8 | 36 | 0.692 | 68.5 |
| 9 | 30 | 0.577 | 64.1 |
| 10 | 34 | 0.654 | 68.1 |

For panels containing up to four loci, *RF*_*norm*_ ranged between 0.62 and 0.89 — far above the threshold of “good concordance” (≤ 0.20) and consistently worse than both top-variable and ML-selected panels of the same size.

Mean bootstrap support remained moderate (57–70%), indicating unstable clades even with a small number of loci.

When panel size was increased to 9–10 loci, RF distances decreased to 0.58–0.65, but these values were still inferior to the ML panels, which achieved the same or lower RF values with higher bootstrap support (e.g., 86.9% at 10 loci).

Thus, selecting loci around the median variability value does not ensure reliable reproduction of the reference topology: topological discordance remained >0.57 even with 10 loci, and bootstrap support did not exceed 70%. These findings demonstrate that moderate variability alone is insufficient for strain-level phylogeny, whereas systematic selection (either by maximum entropy or ML ranking) yields substantially better results at equal or smaller panel sizes.

### 3.6 Limitations of the approach

Despite the promising results of the ML-based selection of informative loci for phylogenetic analysis of *N. gonorrhoeae*, several fundamental limitations of the developed methodology must be noted.

#### Sampling and representativeness

Reference panel size. The use of the WHO-2024 panel of 29 strains ensures high data quality and controlled diversity, but it may not fully reflect the global genetic diversity of *N. gonorrhoeae*. Newly emerging strains may harbor allelic variants absent from this reference collection, which could reduce classification accuracy for novel isolates.

#### Methodological and technical limitations

The accuracy of ortholog detection critically depends on the quality of automatic gene annotation in the source assemblies. Errors in predicted CDS regions or imprecise sequence boundaries can generate artifactual orthogroups and reduce the reliability of phylogenetic reconstruction. The clustering algorithm implemented in OrthoFinder may also incorrectly group paralogous genes or split orthologs with highly variable sequences. In addition, the embedding approach using DNABERT requires substantial computational resources and is constrained by the maximum input length of 512 nucleotides, which may lead to loss of information when analyzing long genes or regions containing multiple insertions/deletions.

#### Biological limitations

All quality metrics (RF distance, bootstrap support) were calculated relative to the reference tree constructed from the same strain panel. Independent validation with newly sequenced isolates of known epidemiological linkage is required to confirm the practical applicability of the method. Moreover, the Robinson–Foulds distance measures only topological differences and does not account for branch lengths, potentially underestimating biologically meaningful distinctions among closely related strains.

### 3.7 Analysis of Antibiotic Resistance Genes

A comprehensive analysis of antibiotic resistance profiles was carried out to identify key genetic variants and assess the prevalence of different resistance mechanisms. The CARD database (v 3.2.5) served as the primary reference, and all computational workflows were implemented in Python (v 3.9.12.

Significant variants were detected in both core genes (e.g., *16S rRNA, PBP1, PBP2, gyrA, parC*) and accessory genes (e.g., *TEM-1, tet(M)*). The heatmap “Presence of Resistance Genes by Strain” (Figure 1) displays a binary panel indicating whether each resistance gene was present or absent in the analyzed strains. Green cells mark the presence of a specific gene, while white cells indicate its absence.

**Figure 1:**
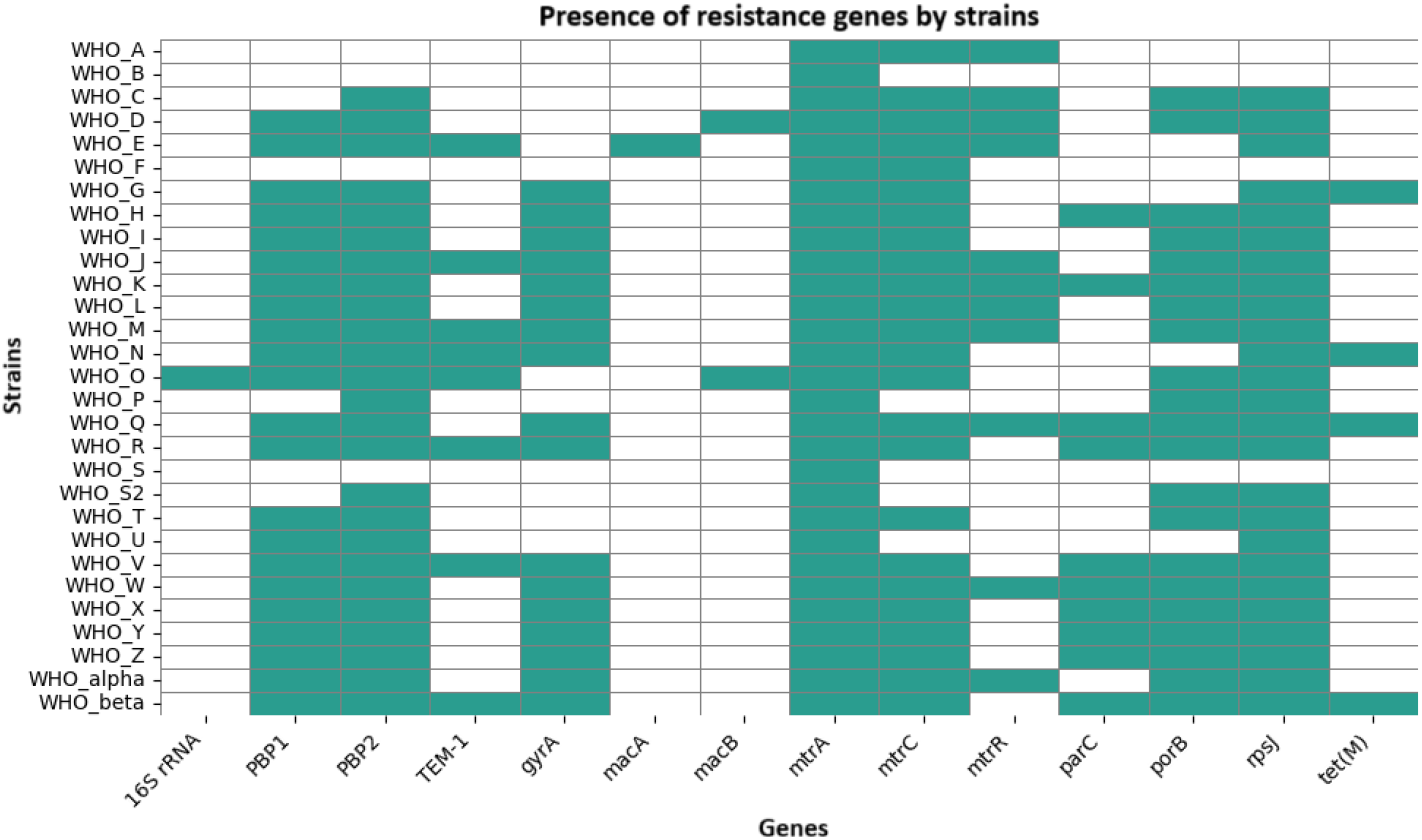
Presence of resistance genes across strains.

The most frequently represented antibiotic classes were *β*-lactamase and macrolides (Figure 2).Among specific resistance-associated genes, *mtrA, penA, rpsJ*, and *mtrC* were the most prevalent.

**Figure 2:**
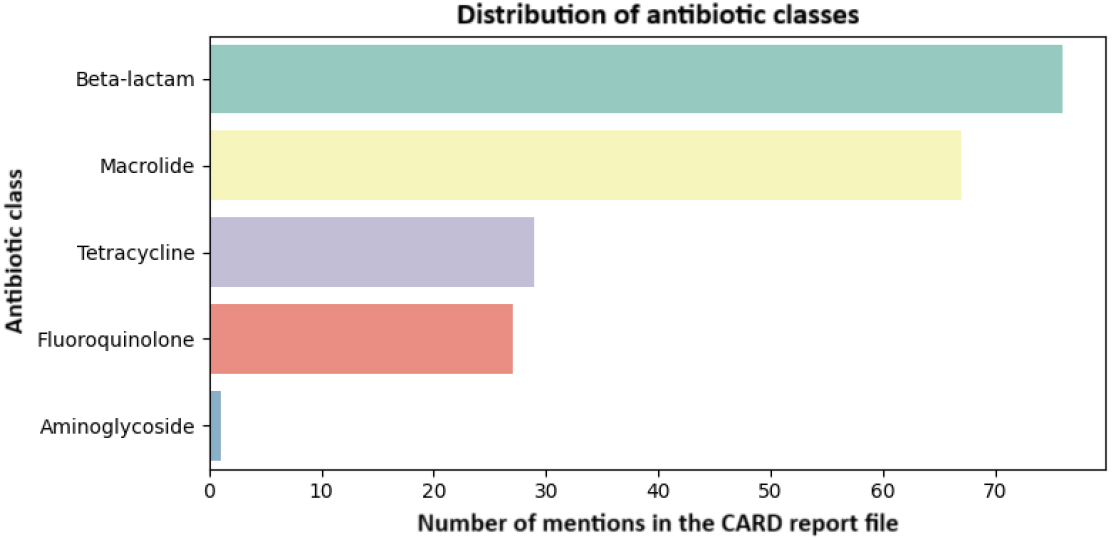
Distribution of antibiotic classes.

The histogram in Figure 3 illustrates the frequency of different genes and their associated resistance mechanisms. Each bar corresponds to a specific gene, and the color coding within the bar indicates the relative contribution of different resistance mechanisms linked to that gene. For example, *mtrA* is predominantly associated with the MtrCDE efflux system, whereas *penA* is largely linked to altered PBP2. This visualization highlights which genes represent the most common determinants of resistance and the mechanisms they primarily mediate.

**Figure 3:**
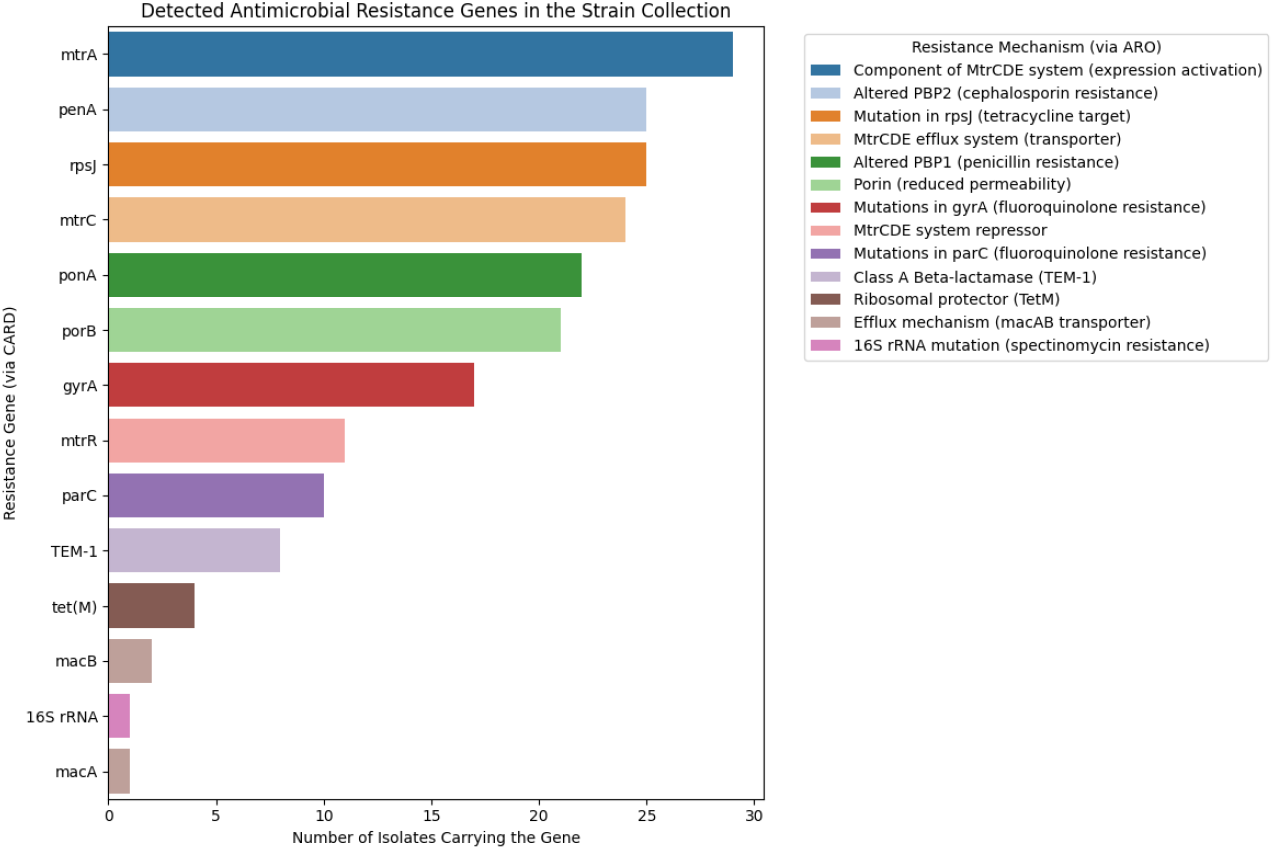
Distribution of genes across strains.

Figure 4 presents a heatmap summarizing the overall resistance patterns of the analyzed strains to different classes of antibiotics. For instance, strains WHO_D, WHO_K, and WHO_N exhibit high levels of resistance to erythromycin, whereas other strains remain susceptible or display lower resistance levels. The histogram in Figure 5 identifies strains with the most extensive resistance profiles. The most enriched strains, each harboring 10 resistance genes, were WHO_Q and WHO_beta.

**Figure 4:**
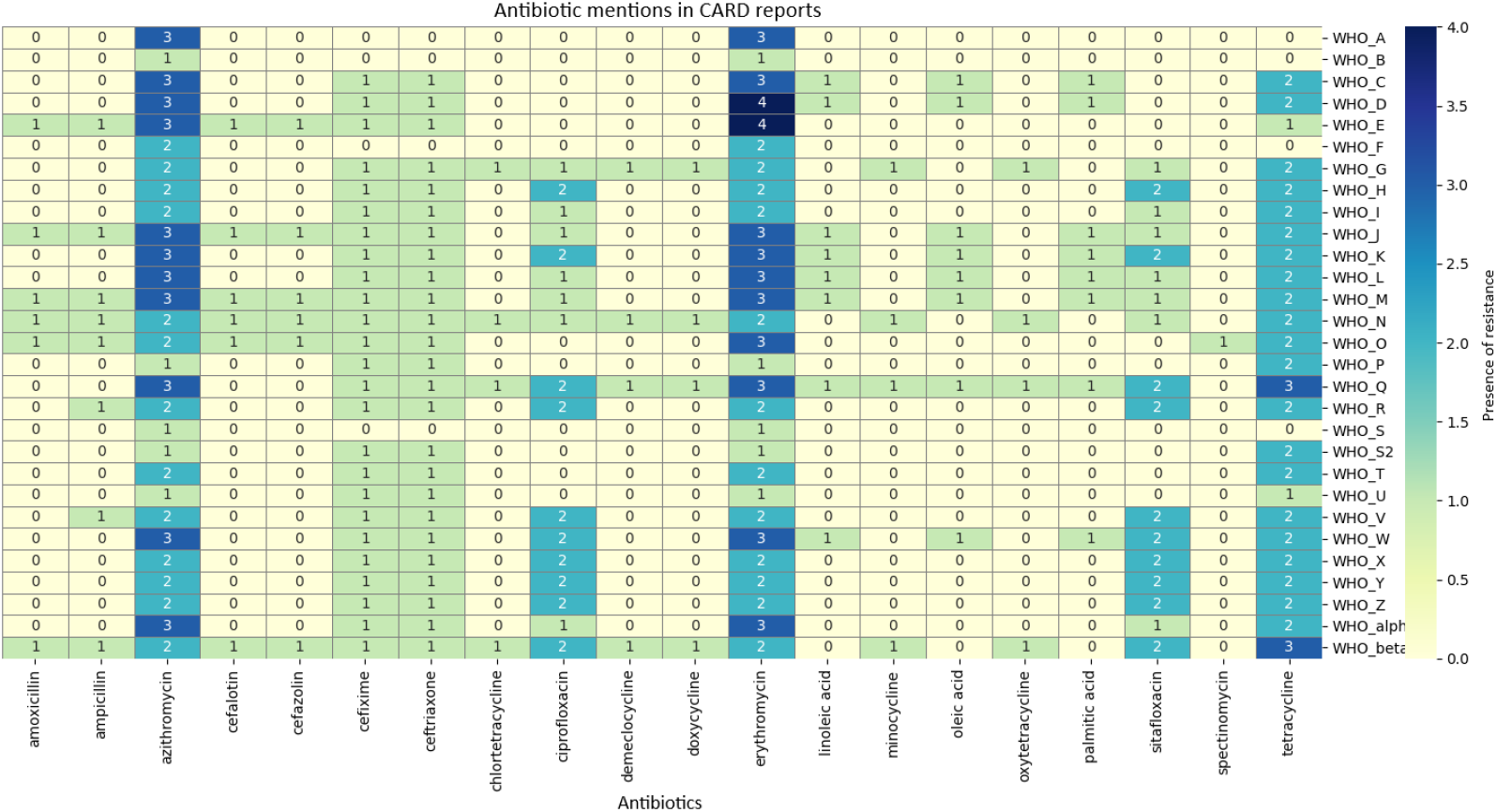
Heatmap of antibiotic mentions in CARD reports.

**Figure 5:**
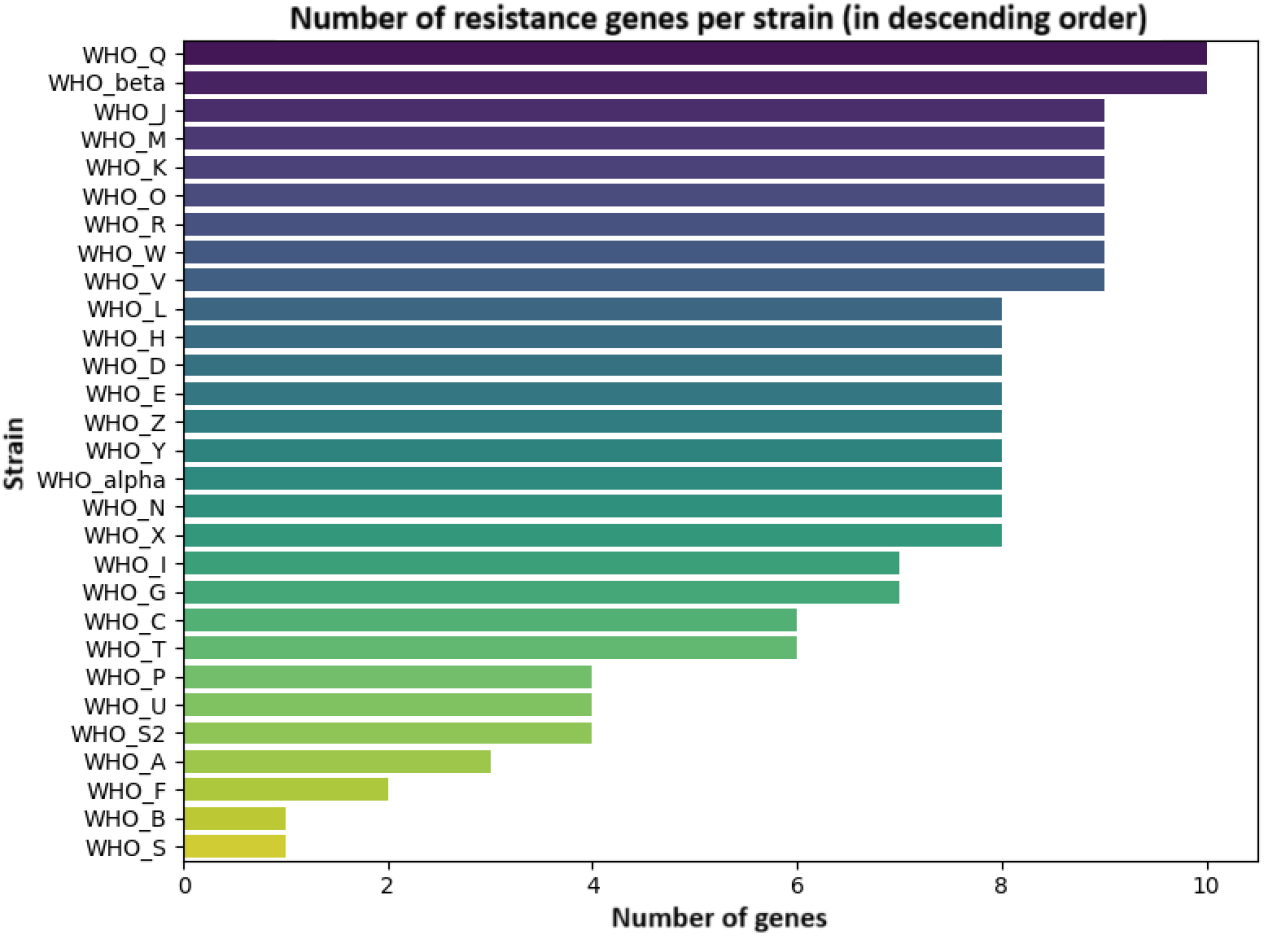
Number of antibiotic resistance variants per strain.

To identify specific mutations contributing to resistance, a detailed analysis of SNP alleles (single-nucleotide polymorphisms) was performed within key genes, including *penA, porB, gyrA, parC, mtrCDE, TEM-1, tet(M)*, and others. The resulting SNP allele dataset provides a resource for further exploration of relevant accessory genes. It should be emphasized that this dataset derives from deep sequencing and is intended for subsequent independent bioinformatic investigation of specific mutations.

The analysis of resistance mechanism frequencies (Figure 6) shows that, for erythromycin and azithromycin, the dominant mechanism is antibiotic efflux.

**Figure 6:**
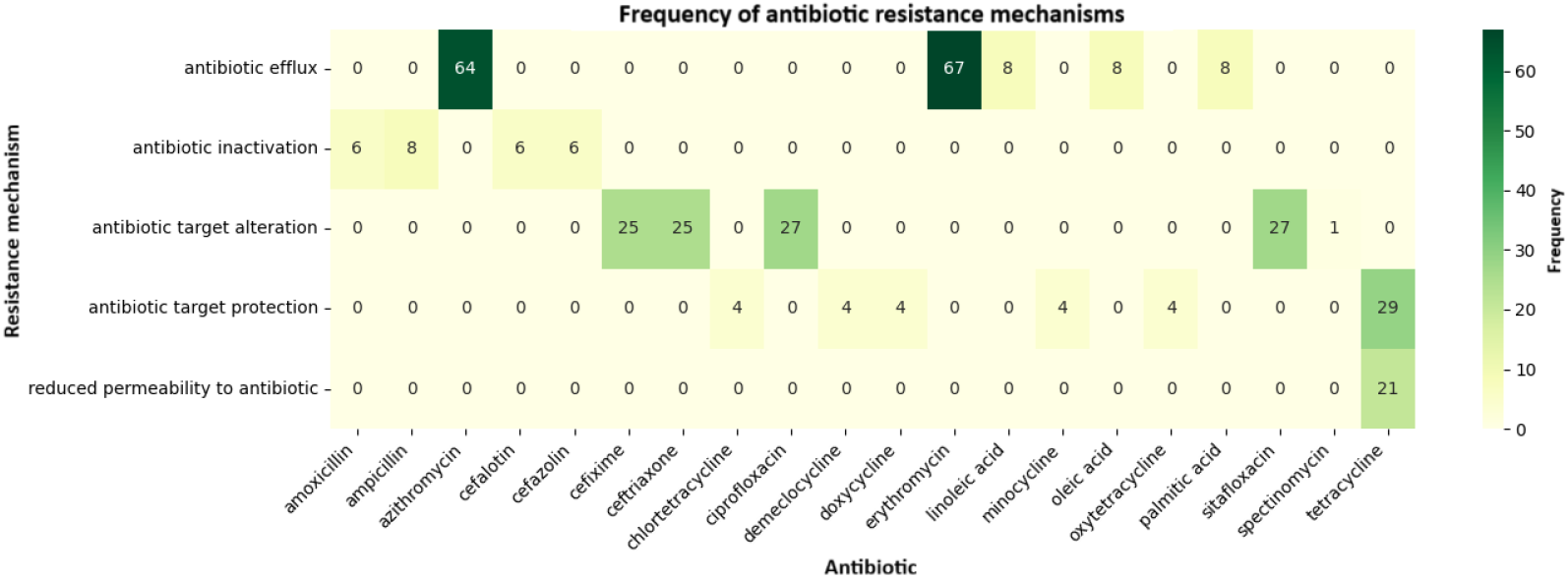
Distribution of resistance mechanisms by antibiotic.

Finally, the chart in Figure 7 demonstrates that the most common combinations involve efflux mechanisms for macrolides and target modification for *β*-lactamase and fluoroquinolones. This histogram offers valuable insights into the predominant strategies employed by bacteria to develop resistance against different antibiotics and may inform the design of more effective treatment strategies and measures to curb the spread of resistance.

**Figure 7:**
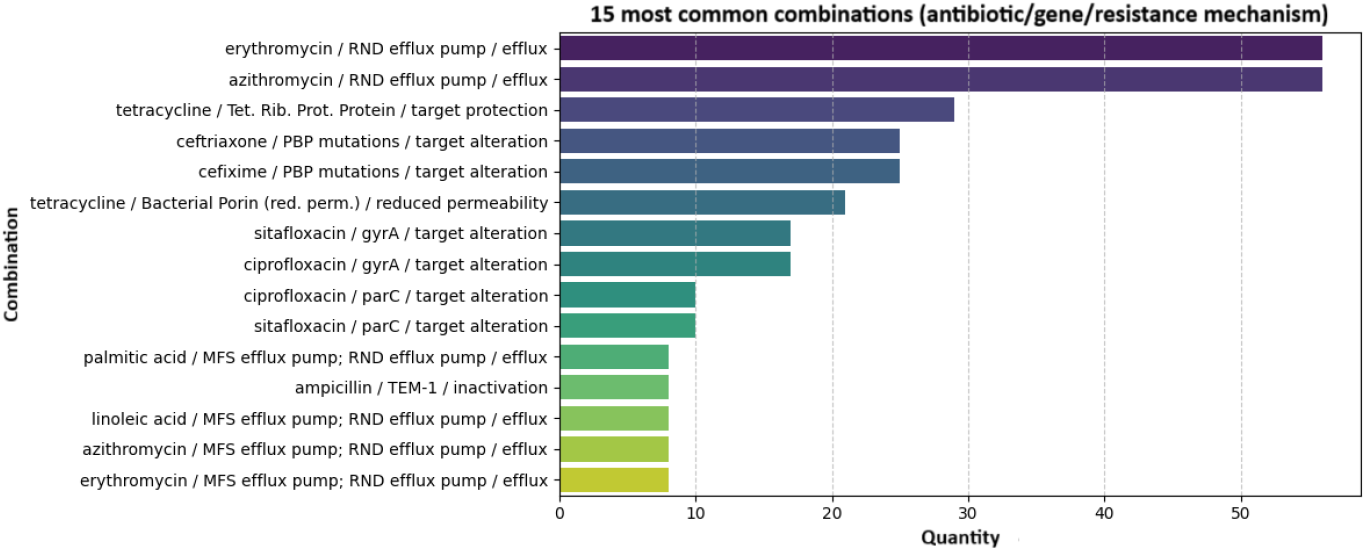
Antibiotic/Gene/Mechanism combinations.

For reproducibility, the complete pipeline, including RGI-parsing and visualization scripts, has been made publicly available in a GitHub repository [25].

### 4 Conclusion

The machine-learning–based approach developed for selecting informative loci demonstrated high efficiency in the phylogenetic analysis of *Neisseria gonorrhoeae*, substantially outperforming classical typing schemes. Machine-learning–selected two-gene panels achieved a normalized RF divergence (*RF*_*norm*_ = 0.769 with a mean bootstrap support of 76.1%), which was markedly better than both panels consisting of the two most variable loci (*RF*_*norm*_ = 0.846, bootstrap = 79.8%) and the classical two-gene NG-MAST scheme (*RF*_*norm*_ = 0.923, bootstrap = 72%). In the most economical configuration of three to four loci, normalized RF divergence decreased to 0.770, and with the expansion to ten genes it reached 0.650; in this setting, mean bootstrap support exceeded 90%, ensuring reliable recovery of strain-level structure.

The successful integration of Random Forest and DNABERT embeddings enables the automated design of molecular typing schemes without the need for full-genome analysis, making the method accessible for routine epidemiological surveillance. The results confirm the feasibility of accurate identification of *N. gonorrhoeae* strains, tracing transmission routes and outbreaks, and monitoring the spread of multidrug-resistant strains, while substantially reducing the costs and labor associated with routine genotyping in clinical laboratories.

Despite these advances, certain limitations remain. Even for the ten-locus panel, the absolute divergence corresponds to roughly 30 out of 52 possible bifurcations. The high rate of recombination in the gonococcal genome produces a mosaic pattern of variation, where loci carrying a “vertical” signal quickly become saturated with parallel substitutions. Moreover, the training dataset of 29 reference strains does not capture the full global diversity, meaning that each new mutation can disproportionately affect the distance matrix. Aggregated Random Forest statistics also do not always distinguish between convergent and genuinely phylogenetic changes.

To address these challenges, a three-stage extension of the methodology is planned. First, the dataset will be expanded with several hundred publicly available genomes, ensuring representation of rare and geographically restricted clusters. Second, the feature space will be enriched with metadata — including isolation date and location, minimum inhibitory concentration phenotype, and resistance determinants — to disentangle signals of selective pressure from vertical evolution. Third, the classical Random Forest is expected to be replaced with more flexible architectures: combinations of gradient boosting, which account for interactions among orthogroups, and graph neural networks capable of leveraging recombination-informative metrics. Such methodological development will further enhance accuracy and reliability across highly diverse strain collections.

## Data availability

The complete pipeline, including scripts, environment configurations, and example data for reproducing analyses, has been deposited at https://github.com/MPHRS/NIRM

## Abbreviations and Definitions

Accessory genes: Genes that can be gained or lost by organisms; not strictly required for survival, but often conferring adaptive advantages such as antibiotic resistance.
Antibiotic resistance: The ability of microorganisms to survive and reproduce in the presence of antibacterial agents that normally inhibit or kill them.
ARG (Antibiotic Resistance Gene): A genetic sequence encoding a protein or RNA that provides resistance to one or more antibiotics.
CARD (Comprehensive Antibiotic Resistance Database): A curated database of antibiotic resistance determinants.
Core genes: Genes present in most or all members of a given group of organisms, typically essential for survival and basic cellular functions.
Core loci: Genes present in all strains of a species.
Efflux: An active mechanism of expelling substances (in this case, antibiotics) from the cell via specialized protein pumps.
HMM (Hidden Markov Model): A statistical model used to represent sequence patterns.
Housekeeping genes: Genes that are consistently expressed in cells, typically essential for fundamental processes.
MAE (Mean Absolute Error): A regression metric measuring average absolute deviation between predicted and observed values.
MFP (ModelFinder Plus): An automated evolutionary model selection algorithm in IQ-TREE.
MLST (Multilocus Sequence Typing): A typing scheme based on seven housekeeping genes.
NG-MAST (Neisseria gonorrhoeae Multi-Antigen Sequence Typing): A two-gene typing scheme (*porB, tbpB*).
NG-STAR (Neisseria gonorrhoeae Sequence Typing for Antimicrobial Resistance): A resistance-associated typing scheme for *N. gonorrhoeae*.
Orthogroups: Groups of orthologous genes that derive from a common ancestral gene through speciation.
Pairwise distance (Hamming): A metric of sequence dissimilarity.
Parsimony-informative sites: Positions in a sequence alignment that provide phylogenetic information.
PBP (Penicillin-Binding Protein): Enzymes involved in bacterial cell wall peptidoglycan synthesis, serving as targets of *β*-lactam antibiotics; mutations in PBPs can confer resistance.
Phylogenetic tree: A graphical representation of evolutionary relationships.
Polytomies: Unresolved nodes in a phylogenetic tree.
RND efflux pump (Resistance–Nodulation–Cell Division): A family of transmembrane proteins forming efflux pumps that actively expel a wide range of antibiotics and other toxic compounds.
Robinson–Foulds distance (RF): A metric for comparing tree topologies.
Shannon entropy: A measure of variability in genetic sequences.
SNP alleles (Single Nucleotide Polymorphism alleles): DNA sequence variants differing by a single nucleotide at a specific position.
Strain clades: Groups of strains sharing a common ancestor, defined through phylogenetic analysis.
Tetracycline Ribosomal Protection Proteins: Proteins that bind to ribosomes and protect them from the action of tetracycline antibiotics.
UFBoot (Ultrafast Bootstrap): A method for assessing branch support in phylogenetic trees.

## Appendix

**Table 6:**
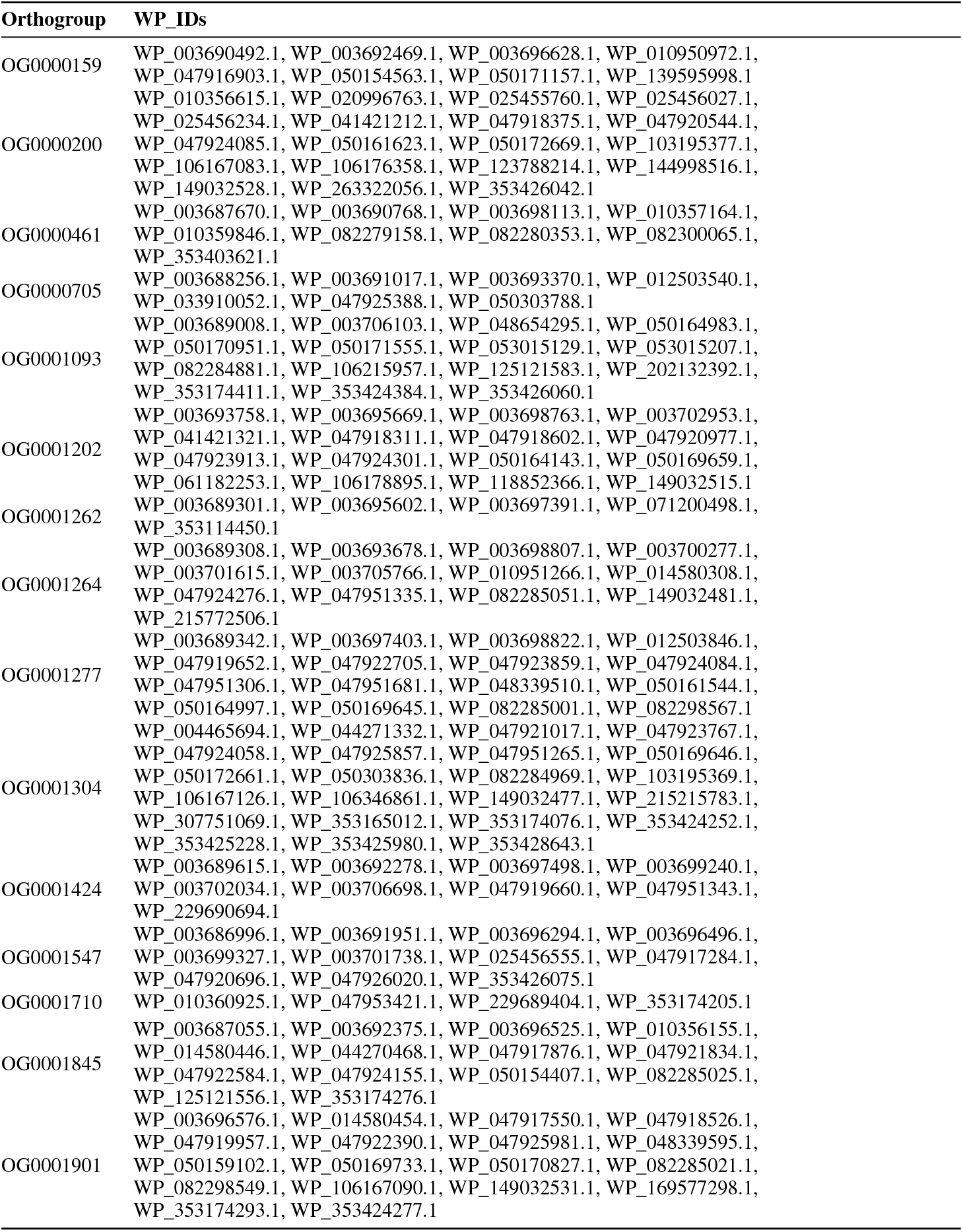
Orthogroups and corresponding WP_IDs.

